## Supplemental figures for "Mutational hotspot in the SARS-CoV-2 Spike protein N-terminal domain conferring immune escape potential"

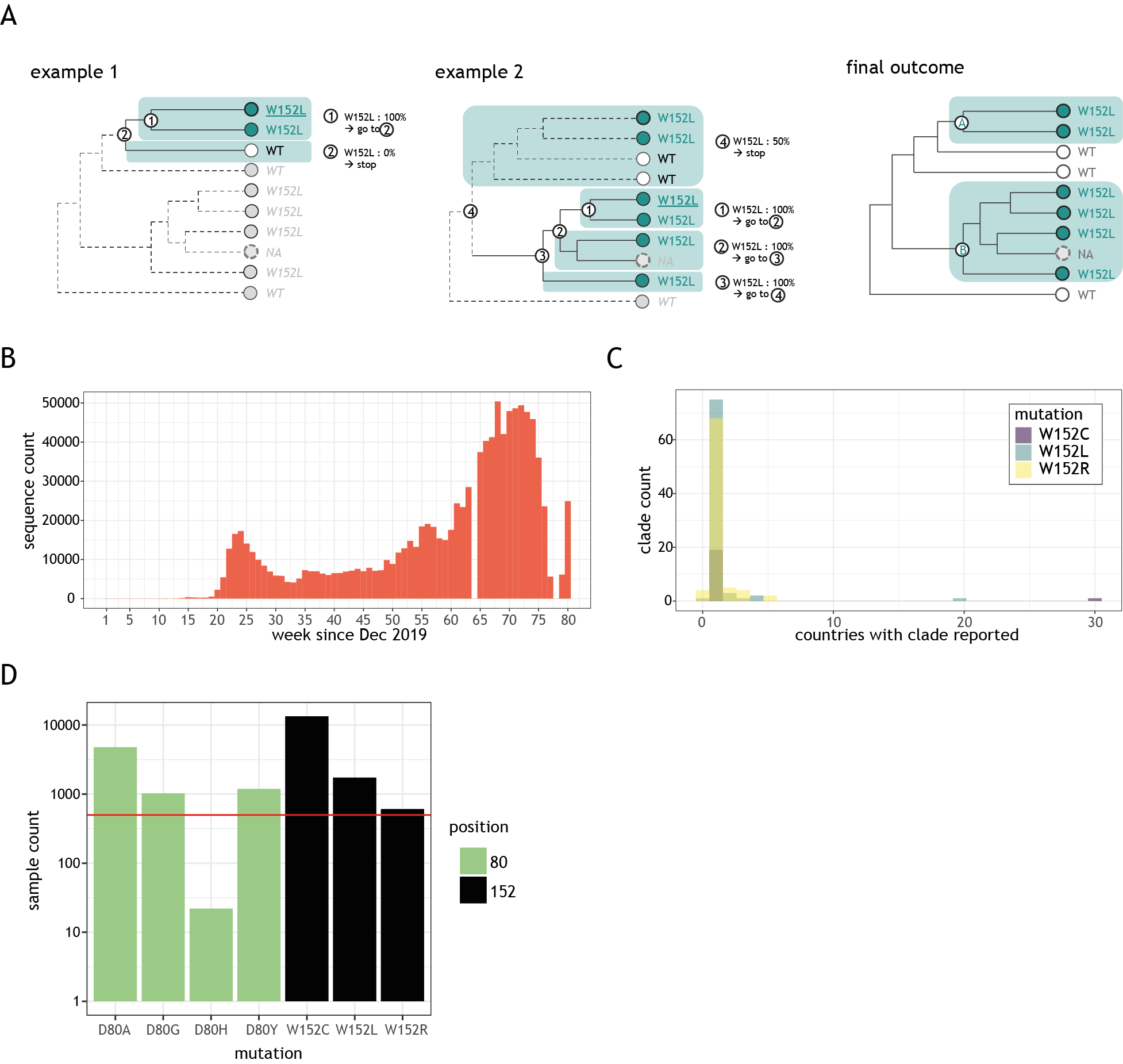

**Figure S1**

(**A**) Heuristic applied to identify and delineate independent W152 clades (see Methods for detailed explanation). (**B**) Number of weekly deposited GISAID sequences (y axis) as a function of collection date (in weeks since December 2019, x axis). (**C**) Count of clades bearing W152 substitutions as a function of the number of countries where clade was reported. (**D**) Count of GISAID sequences reported for diverse substitutions at positions D80 and W152 for which at least 500 cases were reported for at least 3 distinct mutations; red line indicates 500 sequences.

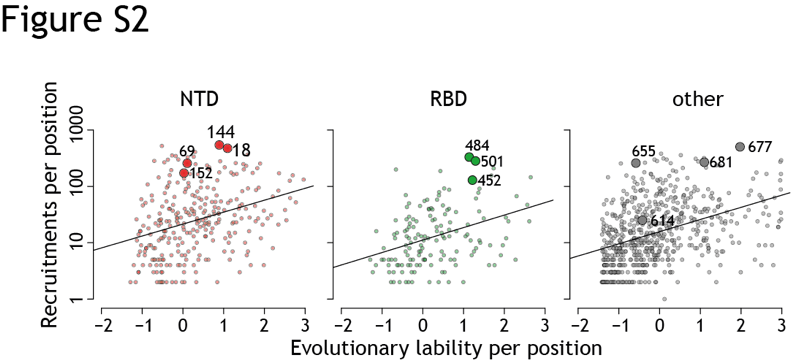

**Figure S2**

Plot showing the relationship between the number of mutation recruitments (y-axis) for each Spike position in NTD (red), RBD (green) or other parts of the protein (grey) and the evolutionary lability score (x-axis) computed for related Coronaviridae; chosen mutation positions are indicated.

The evolutionary lability of Spike residues was estimated using methods made available via the Consurf server ^47^. In this approach homologous entries are fetched from public protein databases, undergo multiple sequence alignment, a phylogeny is built, and the evolutionary rate of each Spike residue is estimated. The obtained rates are then normalized to rank residues from “most conserved” (negative values) to “most labile” (positive values) in the protein of interest. We used the default Consurf parameters (as provided from <https://consurf.tau.ac.il/>) starting with a HMMER search against Uniref90 (<https://www.uniprot.org/help/uniref>) initiated with the Spike amino-acid sequence available in <https://www.rcsb.org/structure/7DDD>. The e-value cut-off was set at 10^-4^ and SARS-CoV-2 entries were manually excluded so as to produce an independent conservation analysis based on 292 representative Coronaviridae sequences.
